# A critical signaling role for diacylglycerol in phagocytosis of *M. tuberculosis*

**DOI:** 10.64898/2026.03.20.713247

**Authors:** Alec Griffith, Michelle Garcia, Gaelen Guzman, Fikadu G. Tafesse

## Abstract

*Mycobacterium tuberculosis* (*Mtb*) establishes infection by entering host phagocytes through phagocytosis. While host lipids are known to influence this process, the specific contribution of the signaling lipid diacylglycerol (DAG) remains poorly defined. Here, we identify DAG as a critical regulator of phagocytosis. Disruption of DAG production, through inhibition or genetic deletion of adipose triglyceride lipase (ATGL) and phospholipase C gamma 2 (PLCγ2), two major pathways that generate cellular DAG pools, markedly reduced uptake of both *Mtb* and zymosan-coated beads. Notably, loss of ATGL or PLCγ2 did not impair receptor trafficking to the cell surface or cargo binding, indicating that DAG is not required for phagocytic recognition or initiation, but instead for a later step in phagosome formation. Mechanistically, cells lacking ATGL or PLCγ2 displayed constitutive phosphoinositide 3-kinase (PI3K) phosphorylation, suggesting that dysregulated intracellular signaling prevents completion of phagocytosis. These findings uncover a previously unappreciated role for DAG biosynthesis in coordinating intracellular signaling required for phagocytosis and provide new insight into host pathways that govern *Mtb* entry.

## Introduction

Phagocytosis is a key process of innate immunity and one of the earliest cellular defenses mounted against intracellular bacterial pathogens^1,2^. This includes *Mycobacterium tuberculosis* (*Mtb*), the causative agent of tuberculosis (TB) and a leading infectious cause of morbidity and mortality worldwide^3^. Phagocytosis involves the recognition of the pathogen, signaling cascades, membrane and cytoskeleton remodeling, and finally pathogen internalization^4^. Professional phagocytes such as macrophages, neutrophils, and dendritic cells are able to expertly take up, kill, and present pathogenic bacteria to induce a larger immune response. Because *Mtb* both depends on and subverts macrophage function to establish infection, defining the molecular signals that control this pathway is essential for understanding TB pathogenesis and identifying host-directed vulnerabilities.

Pathogen uptake by phagocytes begins with recognition of microbes such as *Mtb* through a diverse set of surface receptors such as Fc receptors^5^, toll like receptors^6^, Dectin-1^5^, and others^1,7^. Receptor engagement triggers kinase- and effector-driven signaling cascades that rapidly reorganize the plasma membrane and underlying actin cytoskeleton to promote particle engulfment^1,8^. A central organizing principle in this early phase is local lipid remodeling^9^, classically attributed in large part to phosphoinositide interconversions, where phosphoinositide 3-kinase (PI3K)-dependent conversion of PI(4,5)P₂ to PI(3,4,5)P₃ helps coordinate actin polymerization and membrane extension to form a phagocytic cup^10^, or alternatively supports membrane invagination during “sinking” phagocytosis^11^. These tightly choreographed events culminate in enclosure of the bacterium within a nascent phagosome on the timescale of seconds^12^.

At every step of the phagocytosis process, membrane lipids function not only as structural building blocks but also as spatially and temporally restricted signals that dictate receptor organization, cytoskeletal remodeling, and phagosome maturation. Prior studies highlight, for example, roles for sphingomyelin in phagocytosis of *Candida albicans*^13^ and in receptor clustering during recognition of pathogenic *Mtb*^14^. Among lipid second messengers implicated in phagocytic signaling, phosphoinositides have been comparatively well mapped^1,9,15–18^. In contrast, the contributions of diacylglycerol (DAG), despite its established signaling capacity in many cellular contexts, remain far less defined in the context of phagocytosis.

DAG is structurally simple-two fatty acyl chains esterified to a glycerol backbone and, notably, no polar head group (Fig. 1A). Yet, this minimalist structure supports a potent signaling role at cellular membranes^19–21^. Through its cone-shaped geometry and its ability to recruit and activate DAG-responsive effectors, DAG can reshape local membrane properties and couple receptor engagement to downstream cytoskeletal and trafficking programs^20,22^. Accordingly, both the abundance and spatial distribution of DAG are tightly regulated, with cellular pools supplied by multiple biosynthetic routes. PLC-mediated hydrolysis of PI(4,5)P₂ at the plasma membrane generates DAG and inositol 1,4,5-trisphosphate (IP₃), rapidly producing DAG at sites of immune receptor signaling and actin remodeling^19^. DAG can also be produced through metabolically distinct pathways, including ATGL-mediated hydrolysis of triacylglycerol (TAG) to yield DAG and a free fatty acid^21^, and sphingomyelin synthase (SMS)-dependent sphingomyelin synthesis from phosphatidylcholine and ceramide, which generates DAG as a byproduct^23,24^. A simplified diagram of these pathways is illustrated in Figure 1B. Together, these pathways likely generate distinct DAG pools, raising a key unresolved question in phagocytosis: is DAG required for efficient particle uptake, and if so, which DAG-generating pathway supplies the critical pool needed for the phagocytosis process?

**Figure 1:**
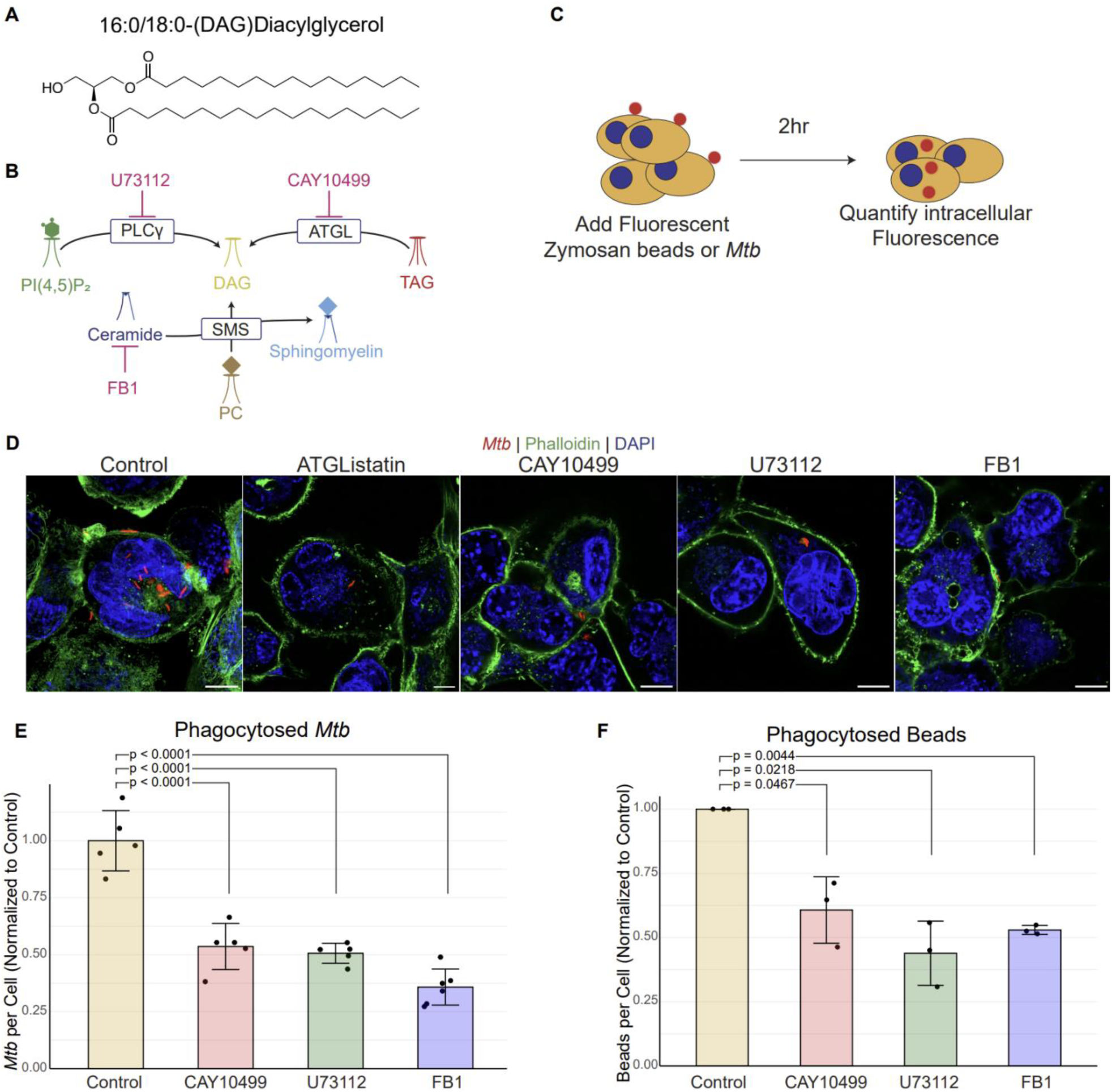
Inhibition of DAG synthesis reduces phagocytic uptake by macrophages. (A) Structure of diacylglycerol (DAG) (B) DAG biosynthetic pathways. (C) Experimental diagram for quantification of phagocytic uptake. (D) Representative images showing phagocytosed *Mtb* in inhibitor treated THP-1 cells. Scale bars represent 10μm. (E) Quantification of uptake of *Mtb* in inhibitor treated cells, normalized to untreated cells. (F) Quantification of uptake of zymosan-coated beads in inhibitor treated cells, normalized to untreated cells. p-value calculated by unpaired t-test with the Benjamini-Hochberg correction.

In this study, we systematically interrogated distinct DAG biosynthetic routes using complementary small-molecule inhibition and targeted genetic perturbations to define the role of DAG in phagocytosis of *Mtb*. We first show that pharmacologic blockade of DAG-generating enzymes reduces phagocytic uptake in macrophages. We then generated macrophage cell lines lacking defined DAG biosynthetic enzymes and found that these genetic perturbations recapitulate the uptake defect. Importantly, the phagocytic defect was not explained by reduced surface receptor expression, indicating a defect in downstream engulfment mechanisms. Finally, we link this failure of uptake to diminished PI3K signaling, consistent with compromised coordination of membrane remodeling and actin dynamics during phagosome formation. Together, our findings build on prior work implicating lipid metabolism in phagocytosis and reveal a previously underappreciated requirement for DAG synthesis in completing this critical innate immune defense mechanism of host cells.

## Results

### Inhibition of DAG synthesis reduces phagocytic uptake by macrophages

We previously showed that SMS activity is important for phagocytosis^14^. Specifically, blocking sphingomyelin (SM) biosynthesis using small-molecule inhibitors, as well as genetic knockout of SMS1-the predominant SMS isoform-significantly reduced macrophage uptake of *Mtb*^14^. Because the SMS-catalyzed reaction generates DAG as a side product, and DAG is a well-established second messenger, it remained unclear whether the observed phagocytic defect reflected loss of SM, loss of DAG, or both. We therefore examined the contribution of cellular DAG to phagocytic signaling. To begin, we treated cultured macrophages with inhibitors targeting DAG synthesis enzymes.

For these studies, we primarily used THP-1 monocytes, a human-derived cell line commonly employed to investigate macrophage phagocytic biology. THP-1 cells were differentiated into macrophage-like cells using phorbol 12-myristate 13-acetate (PMA) and then treated with inhibitors targeting distinct DAG-generating pathways. We used CAY10499 to inhibit ATGL, U73112 to inhibit phospholipase C-γ (PLCγ), and fumonisin B1 (FB1) to reduce ceramide availability (and consequently sphingomyelin production) as a positive control^14^.

To evaluate how DAG biosynthesis influences uptake, we used two complementary cargos. As a BSL-2–compatible surrogate, we employed fluorescent zymosan-coated beads, yeast-derived particles commonly used to quantify phagocytosis without BSL-3 restrictions. To test pathogen-relevant uptake, we also used an mCherry-expressing virulent *Mtb* H37Rv strain under BSL-3 conditions.

Phagocytosis was quantified primarily by microscopy-based assays as described previously (Fig. 1C). Briefly, macrophages were exposed to particles for 2 hours, fixed, and stained with phalloidin to delineate the actin-rich cell boundary, enabling discrimination of internalized cargo (enclosed within the phalloidin-defined perimeter) from extracellular, cell-associated particles (Fig. 1D). Accordingly, inhibitor-treated and vehicle-treated control THP-1 macrophages were incubated with zymosan beads, and internalized particles were quantified and compared across conditions. Blocking DAG biosynthesis by inhibiting ATGL with CAY10499, or inhibiting PLCγ with U73112, each produced a robust and statistically significant reduction in zymosan bead internalization relative to untreated controls (Fig 1F), supporting a requirement for DAG-generating pathways in efficient uptake.

We next asked whether this dependence extends to pathogenic *Mtb*. Inhibiting DAG biosynthesis resulted in an approximately 50% reduction in *Mtb* uptake compared with untreated controls (Fig. 1D-E). As expected, FB1 treatment recapitulated our prior findings, producing a significant reduction in phagocytosis when sphingolipid biosynthesis is blocked. Together, the comparable reduction observed upon inhibiting DAG-generating pathways supports the conclusion that DAG biosynthesis is required for robust, efficient phagocytosis of *Mtb*.

### Generation and Validation of ATGL and PLCγ2 knockouts

Our pharmacological inhibition experiments identified two DAG biosynthetic routes that influence the phagocytic capacity of macrophages, prompting us to validate the contribution of key enzymes in these pathways. We therefore targeted ATGL and PLCγ2 and generated knockout cell lines to directly test how distinct DAG pools regulate phagocytic uptake.

Knockout cells were generated using a CRISPR/Cas9 system with sgRNAs targeting each gene, as previously described^25^. Knockout efficiency was then validated using microscopy-based and biochemical approaches tailored to each target.

Given that ATGL catalyzes the conversion of TAG to DAG on lipid droplets, we assessed ATGL deletion functionally by measuring lipid droplet accumulation^26,27^. Mutant clonal candidates and control cells were loaded with oleic acid and stained with BODIPY 493/503, a neutral-lipid dye that labels lipid droplets. Individual ATGL knockout clones exhibited a doubling of lipid droplet area as measured by BODIPY (Fig. 2A; quantified in Fig. 2B), consistent with impaired TAG mobilization and reduced ability to generate DAG from stored TAG. For PLCγ2, validation was performed by immunoblotting because a reliable commercial antibody is available. Western blot analysis of lysates from the knockout pool showed reduced PLCγ2 protein levels relative to wildtype control (Fig. 2C). We were unable to generate a complete PLCγ2 knockout, likely because PLCγ2 is required for cell survival. Together, these data demonstrate successful generation and validation of ATGL and PLCγ2 knockout macrophages, with clear functional consequences in ATGL-deficient cells and robust depletion of PLCγ2 protein in PLCγ2-deficient cells.

**Figure 2:**
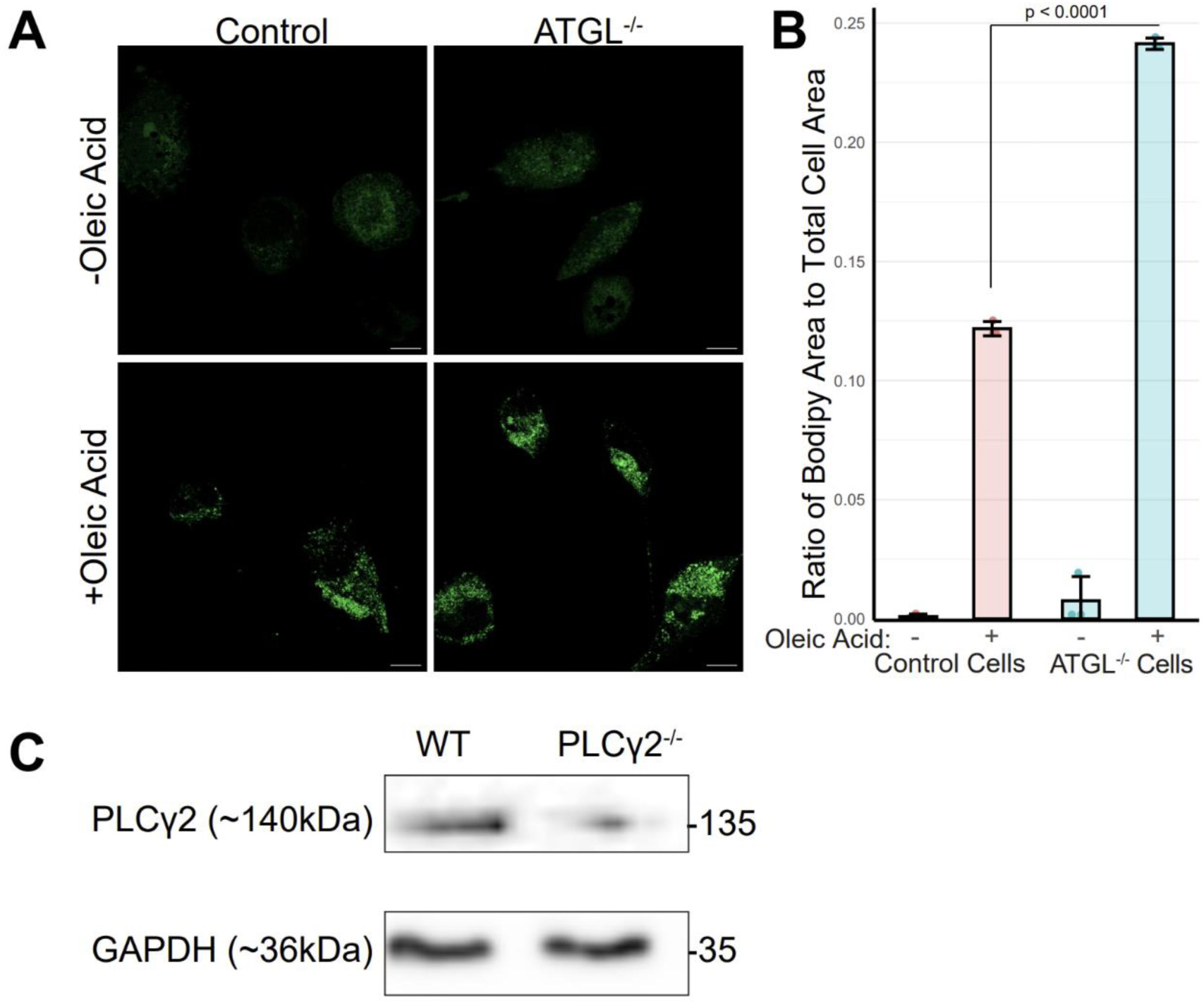
Generation and validation of ATGL and PLCγ2 knockouts. (A) Representative images of BODIPY accumulation in ATGL knockout and wildtype cells. Scale bars represent 10μm. (B) Quantification of images in (A). (C) Western blot demonstrating the knockout of PLCγ2 from pooled samples. p-value calculated by unpaired t-test with the Benjamini-Hochberg correction.

### THP-1s lacking DAG synthesis enzymes have reduced phagocytic uptake

Our inhibitor studies indicated that DAG plays an important role in phagocytosis. To confirm that this effect was due to specific loss of DAG synthesis enzymes rather than off-target effects of the inhibitors, we examined phagocytic function in knockout cell lines. We hypothesized that cells lacking DAG synthesis enzymes would phenocopy the inhibitor-treated cells and display impaired phagocytic uptake.

To test this, we used an experimental setup similar to that shown in Figure 1. THP-1 monocytes were first differentiated into macrophages with PMA and then exposed to either fluorescent zymosan-coated beads or the clinically relevant pathogen *Mtb* as phagocytic cargo. Uptake was quantified by fluorescence microscopy. After incubation with cargo, cells were fixed and stained with phalloidin and DAPI to visualize actin and nuclei, respectively. Internalized beads or *Mtb* were identified as fluorescent cargo enclosed within the phalloidin-defined cell boundary (Figs. 3A,C), and quantification was performed using CellProfiler. Using this approach, we observed nearly a 50% reduction in phagocytic uptake in cells lacking either ATGL or PLCγ2 compared with wild-type cells (Figs. 3B,D). These findings closely mirror the results obtained with pharmacologic inhibition and further support a key role for ATGL and PLCγ2 in phagocytosis.

**Figure 3:**
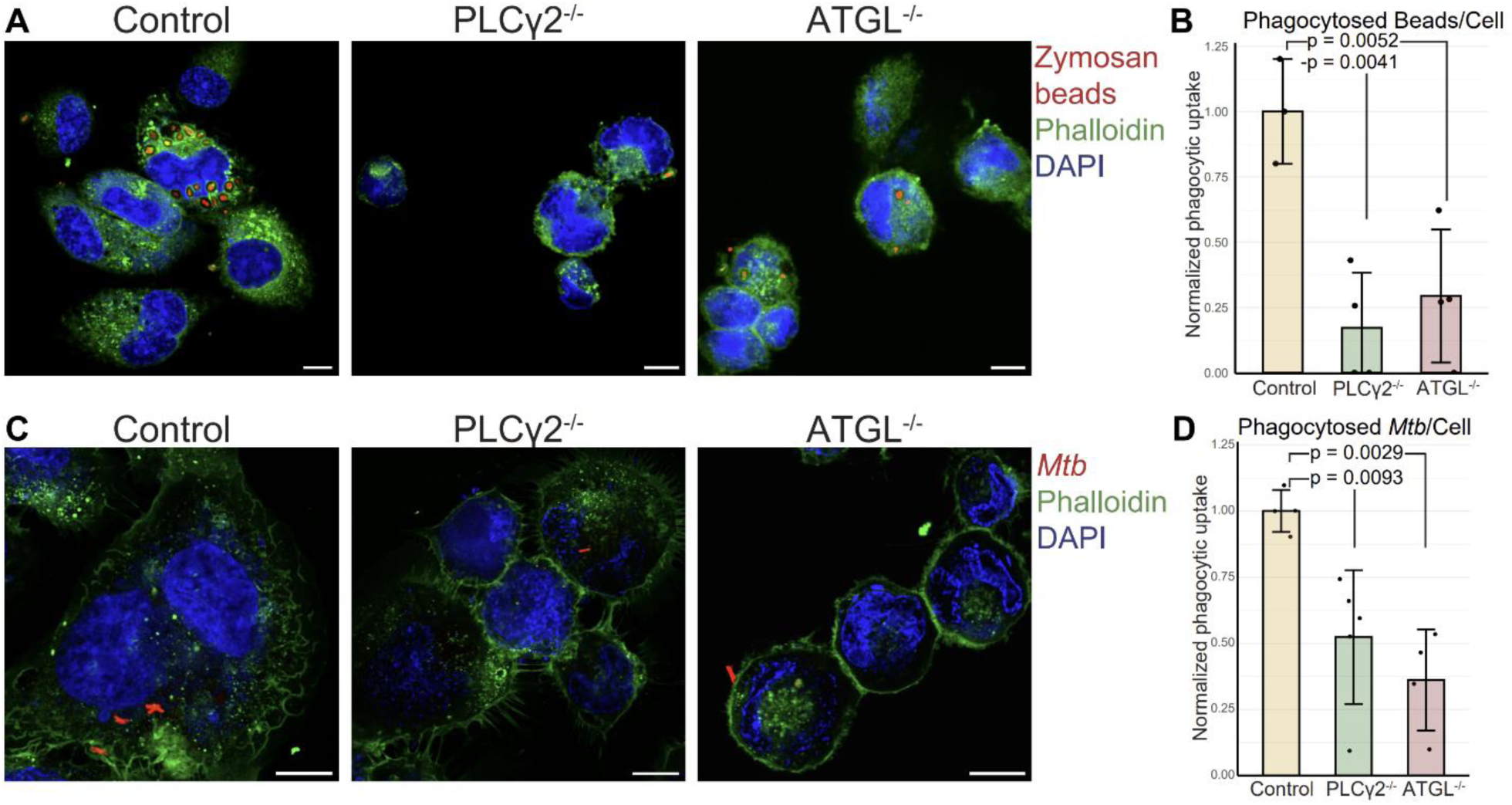
THP-1s lacking DAG synthesis enzymes have reduced phagocytic uptake. (A) Representative images showing phagocytic uptake of zymosan-coated fluorescent beads in control and knockout cells. Scale bars represent 10μm. (B) Quantification of (A). (C) Representative images showing phagocytic uptake of fluorescent *Mtb* in control and knockout cells. Scale bars represent 10μm. (D) Quantification of (C). p-value calculated by unpaired t-test with the Benjamini-Hochberg correction.

### Cargo binding is not impacted in ATGL and PLCγ2 knockout cells

After establishing that ATGL and PLCγ2 are required for cargo uptake, we next sought to define which step of phagocytosis is impaired by reduced DAG synthesis. Because cargo binding and recognition represent the earliest steps of phagocytosis, we first tested whether the observed uptake defect reflected impaired cargo binding. Based on the established role of DAG as a lipid second messenger, we hypothesized that DAG synthesis instead functions at a downstream signaling step following cargo engagement.

To test whether cargo binding is impaired in ATGL- and PLCγ2-mutant cells, we used an assay previously established in our laboratory^28^. Briefly, THP-1 monocytes were differentiated into macrophages with PMA, then cooled to 10 °C to slow cellular processes and prevent progression of phagocytosis. Fluorescent zymosan-coated beads were added, and the cultures were briefly centrifuged to promote contact between beads and cells. Samples were then briefly warmed to 37 °C before washing, fixation, and staining. Fluorescently labeled concanavalin A was used to outline the cell surface, and cells were imaged by confocal fluorescence microscopy (Fig. 4A). Zymosan binding was quantified by counting fluorescent beads associated with the cell periphery while excluding beads that were internalized or not cell-associated (Fig. 4B). Comparison of knockout and wild-type cells revealed no significant difference in cargo binding. These results indicate that loss of ATGL or PLCγ2 does not impair cargo recognition or adhesion, and instead suggest that the reduction in uptake reflects a defect in phagocytic internalization.

**Figure 4:**
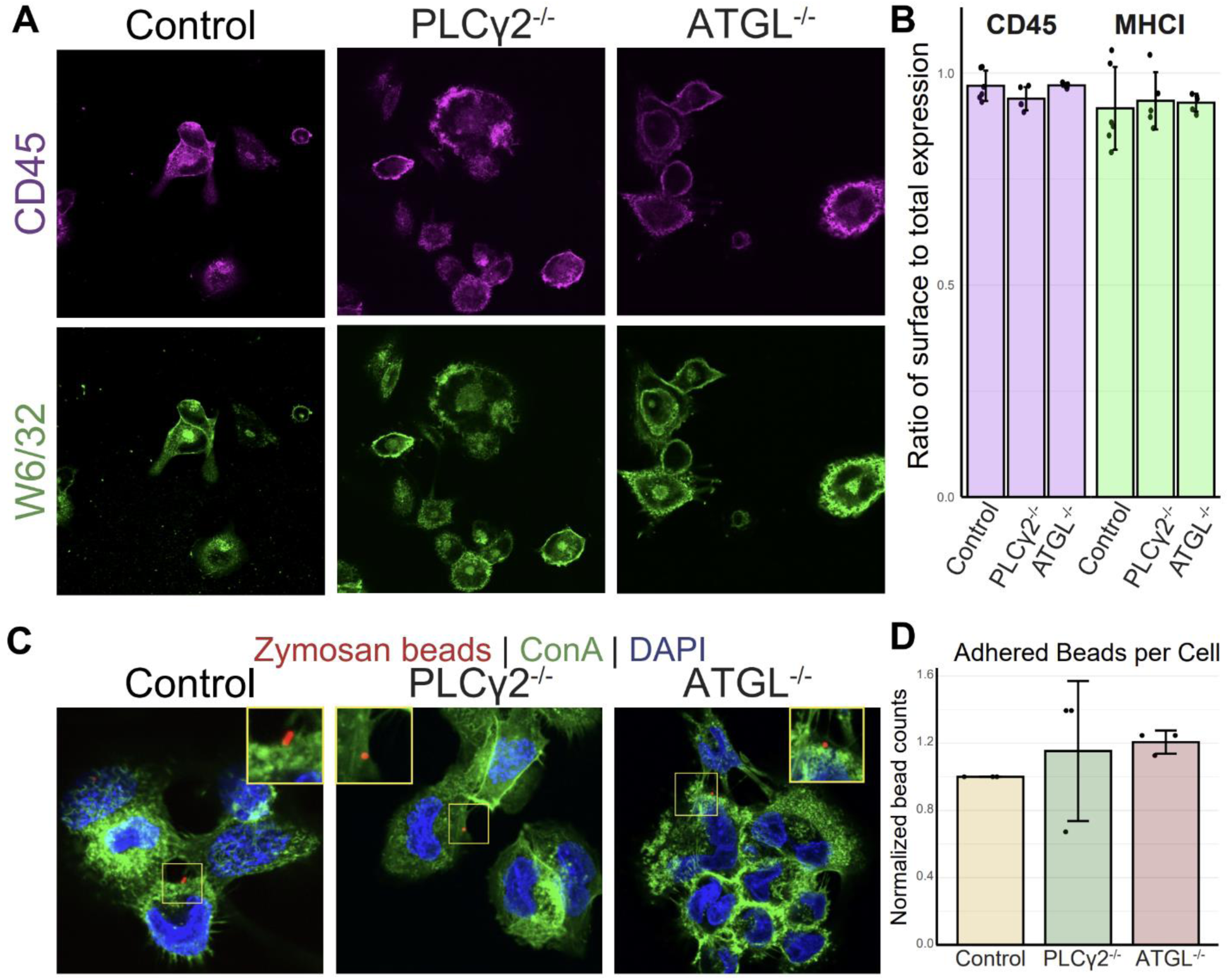
Changes in phagocytic uptake unrelated to cargo binding or receptor expression. (A) Images showing adherence of zymosan-coated fluorescent beads to control or knockout cells. (B) Quantification of (A). (C) Images of permeabilized control and knockout cells showing total expression of CD45 and MHC-I. (D) Quantification of (C). p-value calculated by unpaired t-test with the Benjamini-Hochberg correction.

### Expression and trafficking of surface receptors is unchanged after DAG synthesis knockouts

Having established that knockout cells bind zymosan-coated beads at rates comparable to wild-type cells, we next asked whether receptor trafficking to the cell surface was also intact. Reduced surface expression of key phagocytic receptors could provide an alternative explanation for the impaired phagocytosis observed in the knockout cells.

To measure the surface expression of these receptors, a microscopy-based assay was employed. PMA differentiated wild-type or knockout THP-1 cells were fixed prior to staining. These cells were split into two groups, one was permeabilized, to measure total receptor amount, and the other was left with an intact plasma membrane to measure only surface expression. These cells were then stained with antibodies against CD45, a known surface receptor on immune cells, and class I MHC, a receptor trafficked from the ER to the cell surface (Fig 4C). The fluorescent intensity of these stains was measured and a ratio of intensities between the unpermeabilized and permeabilized samples was calculated. A cell with trafficking defects would have a significantly higher amount of intracellular receptor which would show up as a lower ratio than a cell with a competent trafficking system. Across all tested cell types, no defect was seen in receptor trafficking or overall expression of the tested receptors (Fig 4D). This indicates that cells lacking DAG biosynthesis pathways are able to properly express and traffic receptors to their surface for recognition of cargo to be phagocytosed. This indicates that a defect in a cellular process involving the internalization of cargo is the primary driver of the phenotypes observed in these knockout cells.

### Constitutive PI3K activation induced by perturbations in DAG synthesis

After establishing that ATGL- and PLCγ2-deficient cells have impaired phagocytic capacity, and that this defect is not due to altered cargo binding or receptor surface expression, we next sought to define the underlying mechanism. Given the central role of phosphoinositide 3-kinase (PI3K) signaling in phagocytosis, we examined PI3K phosphorylation following cargo exposure^10^. We hypothesized that loss of DAG synthesis enzymes leads to dysregulated PI3K signaling.

To assess this, we stimulated cells with zymosan-coated beads for 5, 15, 30, and 60 minutes and measured total and phosphorylated PI3K in whole-cell lysates by Western blotting. Total PI3K levels were comparable between wild-type and ATGL- or PLCγ2-deficient cells at all time points following zymosan stimulation (Fig. 5A). In contrast, PI3K phosphorylation differed markedly. In wild-type cells, zymosan induced rapid PI3K phosphorylation that declined after 30 minutes, as expected. However, in both ATGL- and PLCγ2-deficient cells, PI3K phosphorylation remained elevated from baseline through 60 minutes of stimulation, with no evident decline even at the latest time point (Figs. 5B,C).

**Figure 5:**
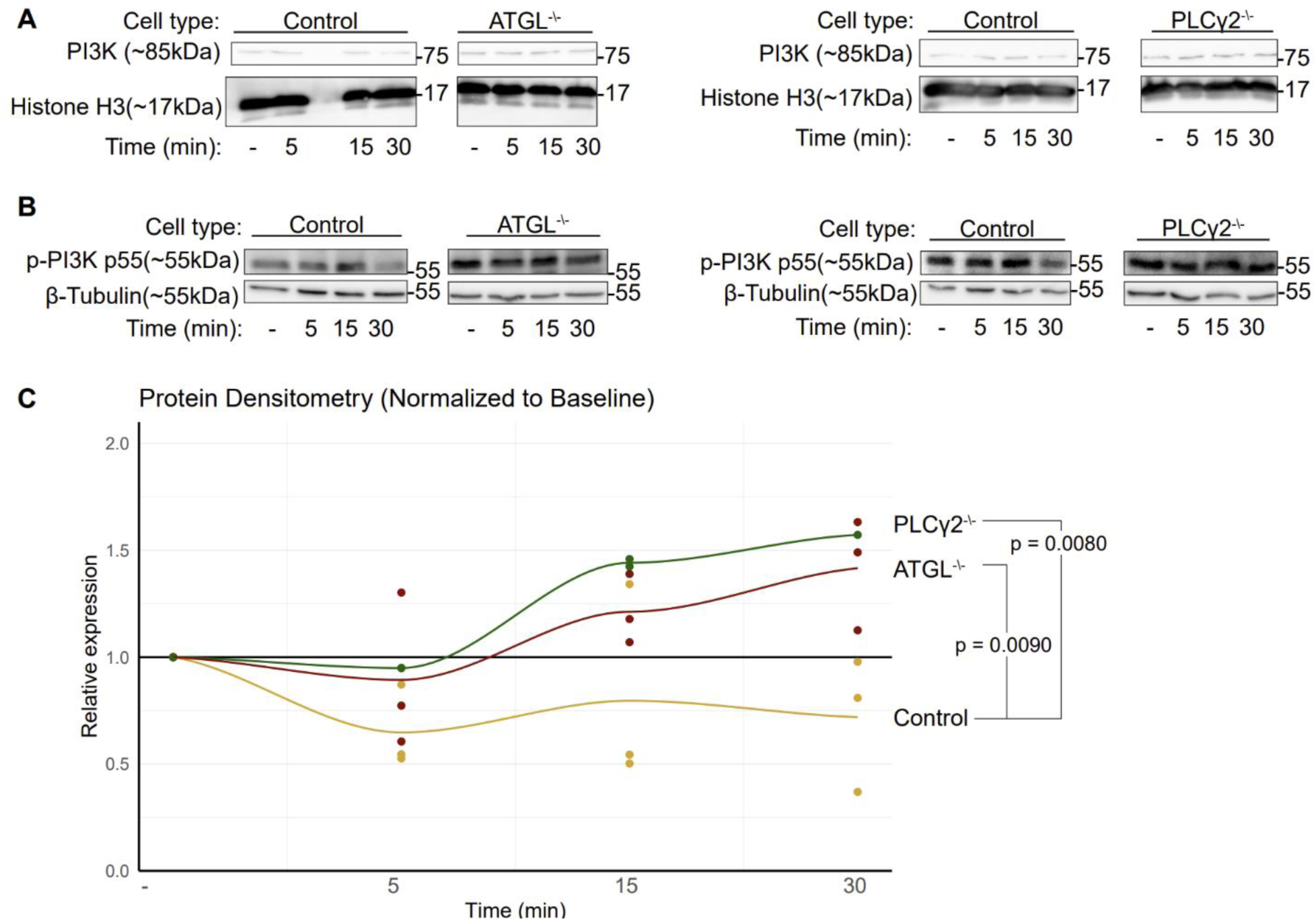
Aberrant PI3K activation induced by perturbations in DAG synthesis. (A) Western blots showing total PI3K expression levels in knockout cells at indicated bead exposure times. (B) Changes in PI3K phosphorylation after indicated exposure times to zymosan-coated beads. (C) Quantification of p-PI3K levels in tested cells. p-value calculated by ANOVA with Tukey HSD post-hoc test.

These data demonstrate that DAG synthesis through the ATGL and PLCγ2 pathways is essential for proper PI3K signaling and efficient completion of phagocytosis. ATGL- and PLCγ2-deficient cells, which are impaired in DAG synthesis, exhibited constitutively elevated PI3K phosphorylation both at baseline and after stimulation with zymosan-coated beads. Given the central role of PI3K and its downstream products in phagocytosis, this persistent phosphorylation likely impairs phagocytic completion. Together, these findings support a model in which DAG synthesis regulates phagocytosis through control of PI3K-dependent signaling.

## Discussion

In this study, we identify DAG as an essential regulator of macrophage phagocytosis and *Mtb* uptake. Genetic disruption of the DAG synthesis enzymes ATGL and PLCγ2 impaired cellular DAG production and significantly reduced phagocytic internalization. Cells lacking ATGL or PLCγ2 showed reduced cargo internalization, but they displayed no defect in cargo binding or cell-surface receptor expression, indicating that DAG synthesis is dispensable for phagocytic recognition but essential for later stages of uptake. We further show that these cells exhibit aberrant PI3K signaling, which likely contributes to the phagocytic defect. Together, our findings establish DAG synthesis as a critical pathway controlling phagocytic completion during *Mtb* entry into macrophages.

DAG is an important molecule for the transduction of cellular signals through not only through the products of its synthesis, but also through its recruitment of adapter proteins. One of the best studied examples of this is the recruitment of protein kinase C (PKC) which has a C1 domain for binding DAG in membranes^20,31^. PKC recruitment to the plasma membrane can lead to the activation of protein phosphatase 2 (PP2) which can negatively regulate the activity of enzymes such as PI3K^32,33^. Dephosphorylated PI3K is unable to convert PIP_2_ to PIP_3_^34^. This conversion is essential for the completion of phagocytosis, PIP_2_ is synthesized at the initiation site of the phagocytic cup and is converted, by PI3K, to PIP_3_ as the phagocytic cup matures and engulfment begins^35,36^. PIP3 is an essential molecule for the completion of phagocytosis through modulation of actin dynamics and maturation of the early phagosome^9,15,16^. It is likely that cells lacking DAG are diminished in their ability to recruit PKC, leading to increased activation of PI3K and subsequent inability to complete phagocytosis.

This study further characterizes the importance of two DAG synthesis enzymes in phagocytosis; ATGL^37^ and PLCγ2^38,39^. Here we demonstrate that these enzymes are required for the completion of phagocytosis of zymosan-coated beads and *Mtb*. This study validates a role for PLCγ2 in plasma membrane signaling and outlines a surprising role for ATGL-produced DAG in regulating phagocytic signaling.

Phagocytosis is an essential component of innate immunity and the clearance of extracellular and intracellular debris^4,40^. It is an essential first step in the process of antigen presentation that leads to the development of adaptive immunity^41,42^. The work discussed here shows new findings that help to further clarify the cellular processes that compose phagocytosis. This work also highlights a previously undescribed role for DAG in this process.

The data demonstrated here demonstrate that DAG is an essential lipid for phagocytic uptake by cells. However, these data merely demonstrate the phenomenon, and more work needs to be done to determine the specific mechanisms by which DAG influences phagocytosis. Specifically, the data here demonstrates that DAG synthesized by both ATGL and PLCγ2 is important for phagocytosis, but that these sources of DAG impact phagocytic uptake differentially. Determining the roles of DAG from different sources will require further study. In addition, the role of the lipid itself is not well characterized in phagocytosis. It remains possible that other functions of ATGL and PLCγ2 are responsible for the phenotype which remain confounded in inhibitor and knockout experiments. Experiments looking at DAG localization or supplementation in these knockouts will be valuable for future studies.

All together, we have demonstrated that the phagocytic ability of macrophages is linked to their ability to synthesize DAG. By generating knockouts of ATGL and PLCγ2 we have demonstrated that these two DAG synthesis enzymes play important roles in the completion of phagocytosis with no impact on surface receptor expression. Finally we have demonstrated that perturbations in DAG synthesis lead to aberrant PI3K signaling downstream of cargo recognition in a manner that differs between ATGL and PLCγ2 knockout cells.

## Methods

### Phagocytosis assays

For experiments making use of zymosan-coated fluorescent beads, cells were plated in 96 well plates at 1 x 10^4^ cells per well or on #1 coverslips in 24 well plates at 5 x 10^4^ cells per coverslip. Cells were then activated with 50nM phorbol 12-myristate 13-acetate (PMA; Sigma-Aldrich, product P8139-1MG) for 48 hours. For phagocytosis experiments using inhibitors, inhibitors were added at their respective concentrations for 72 hours. After activation Alexa Fluor 594-conjugated Invitrogen™ zymosan A (*Saccharomyces cerevisiae*) BioParticles (Fisher Scientific, product Z23374) were added at an MOI of 5 and spun at 175 x g for 3 minutes to force contact between the beads and cells. Plates were then transferred into a 37°C incubator with 5% CO_2_ for 2 hours. After incubation cells were removed and washed 3 times with cold PBS to remove any extracellular beads and then fixed in 4% formaldehyde for 30 minutes.

For experiments making use of dsRed-*Mtb* an MOI of 10 was used.

Coverslips or plates were stained with 0.75 Units/mL of Dylight^TM^ 488 Phalloidin (Cell Signaling Technology, product 12935S) and 2 µg/mL of 4’,6-diamidino-2-phenylindole (DAPI) (Thermo Fisher, product D3571) in PBS containing 5% Bovine Serum Albumin (BSA; Sigma Aldrich, product A7906-100G) and 0.01% Triton-X 100 for 30 minutes. Staining solution was removed and cells were treated with 4% paraformaldehyde (Electron Microscopy Sciences, product 15710) for 30 minutes. 96 well plates were washed 2 x with PBS before imaging and coverslips were rinsed with deionized water and mounted with ProLong Glass Antifade (Thermo Fisher, product P36980). For quantification (Figs 1E-F, 3B,D) images were acquired on a Keyence BZ-X 700 with a Nikon CFI S Plan Fluor ELWD ADM 40XC objective. Images were acquired in a 5 x 5 square and stitched together prior to analysis in CellProfiler software^43^. DAPI was initially used to identify all cell objects as “primary objects” and the bounds of the cell were defined as “secondary objects” using the phalloidin signal. These objects were used to generate a mask which was applied to the bead image allowing for the identification of only intracellular objects.

For visualization (Figs 1D, 3A,C) images were acquired on a laser scanning confocal microscope (Zeiss LSM 900) with a Plan-Apochromat 63x/1.4 Oil DIC M27 objective (Zeiss, product 420782-9900-000) and 405 nm, 488 nm, and 561 nm lasers. These images were processed in FIJI^44^ prior to publication to adjust brightness and contrast.

### Culturing of mammalian and bacterial cells

THP-1 monocytes (ATCC, product TIB-202) were cultured in RPMI 1640 medium (Thermo Fisher Scientific, product 11875119) supplemented with 10% FBS (Sigma-Aldrich, product F0926), 1X Penicillin-Streptomycin (Thermo Fischer Scientific, product 15-140-163), and 1X Non-Essential Amino Acids (Thermo Fischer Scientific, product 11140050). Cells were cultured at 37°C and 5% CO_2_. Prior to activation cells were counted with a hemocytometer using Trypan Blue as counterstain.

All infections with *Mycobacterium tuberculosis* were performed in a dedicated biosafety level 3 laboratory following approved safety guidelines.

The *Mtb* strain H37Rv-dsRed (a gift from the lab of Dr. Dave Lewinsohn) was cultured in Difco Middlebrook 7H9 (Becton Dickinson, product 271310) supplemented with 10% Difco Middlebrook oleic acid-albumin-dextrose-catalase enrichment (OADC; Beckton Dickinson, product L12240), and 0.05% Tween-80 (Thermo Fisher Scientific, product BP337-500). When the 600 nm optical density (OD_600_) of a culture reached between 0.5 and 1.0 bacteria were collected for infection. Prior to infection, bacteria were spun at 2000 x g for 5 minutes, washed with 10mL PBS, and resuspended in RPMI 1640 supplemented as above but without antibiotics. Bacteria were resuspended by passing through a blunt-tipped 18-gauge needle 25 times and passage through a 5 μm filter to break up or remove any clumps. The OD_600_ of this suspension was measured and the concentration of bacteria was estimated using a modified McFarland standard^45^ of OD_600_ = 1.0 = 2.0 x 10^8^ CFU / mL. Further dilutions, as required to reach desired MOI, were performed in RPMI without antibiotics.

### Inhibitor treatment

For experiments where making use of chemical inhibitors previously published concentrations were used. Inhibitors were suspended in DMSO. All inhibitor treatments were conducted for 72 hours after activation of THP-1s. To inhibit DAG synthesis through ATGL CAY10499^46^ (Cayman Chemical, product 10007875) was used at 10 μM. To inhibit PLCγ2 U73112^39,47^ (Cayman Chemical, product 70740) was used at 5 μM. FB1 (Cayman Chemical, product 62580) was used to reduce cellular ceramides to decrease SMS activity at a concentration of 5 μM^14,48^.

### Generation of knockouts

ATGL and PLCγ2 knockouts were generated using the lentiCRISPRv2 plasmid and protocol developed by the Zhang lab^25^. The empty lentiCRISPRv2 plasmid was digested with FastDigest *BsmBI* (New England Biolabs, product R0739S) and dephosphorylated with rSAP (New England Biolabs, product M0371S) overnight at 37 °C. Digested plasmids were purified by gel electrophoresis and extraction with a Monarch gel extraction kit (New England Biolabs, product T1020). Complementary DNA oligonucleotides for the gRNA sequences were synthesized by Azenta and were annealed and phosphorylated using T4 PNK (New England Biolabs, product M0236S) and T4 ligation buffer (New England Biolabs, product B0202S)^49^. The digested plasmid and annealed oligos were ligated overnight at room temperature in the presence of T4 ligase (New England Biolabs, product M0202S) and T4 ligation buffer. This ligated plasmid was transformed into Stbl3 bacteria (Invitrogen, product C7373-03) following the instructions of the manufacturer.

Plasmids were extracted from bacterial cultures using Omega E.Z.N.Z Plasmid DNA Mini Kit miniprep kit (Omega Bio-Tek, product D6945-02) following manufacturer instructions. Replication deficient lentivirus was produced in HEK 293T cells (ATCC, product CRL-3216) by transfection of the complete lentiCRISPRv2 plasmid, psPAX (a gift from Didier Trono, Addgene plasmid #12260), and pMD2.G (a gift from Didier Trono, Addgene plasmid #12259) via Lipofectamine 3000 (ThermoFisher, product L3000015) following manufacturer instructions. After 48 hours of lentiviral production, the supernatant from transfected cells was filtered through a 0.45 micron filter to purify the lentivirus and THP-1 cells were infected overnight. After infection, cells were selected with 2 μg/mL of puromycin (Sigma-Aldrich, product P7255) to select for successfully transduced cells.

### ATGL knockout validation

5 x 10^4^ wild-type or ATGL^-/-^ THP-1 cells were activated with 50 nM PMA for 48 hours on coverslips in 24 well plates. Cells were then treated with 30 μM oleic acid (OA; Cayman Chemical, product 90260) or DMSO vehicle overnight. Cells were washed 3 times with PBS before the addition of BODIPY 505/515 (Cayman Chemical, product 25893) to all cells at 2 μM in PBS for 20 minutes following previously published protocols^26^. Dye was washed off with PBS before cells were fixed with 4% formaldehyde for 30 minutes. Coverslips were stained with DAPI for 10 minutes and mounted with ProLong glass antifade. For quantification (Fig 2 A) images were acquired on a Keyence BZ-X 700 with a Nikon CFI S Plan Fluor ELWD ADM 40XC objective. Images were acquired in a 5 x 5 square and stitched together prior to analysis in FIJI software. Cell and BODIPY areas were measured using thresholding and the Analyze Particles module. The ratio of BODIPY area to cell area was calculated.

### Antibodies used

For protocols with primary and secondary antibody staining, the following antibodies were used at indicated concentrations. Anti-PLC gamma2 (1:1000; Cell Signaling Technologies, product 3872T), anti-GAPDH (Cell Signaling Technologies, product 8884S), Alexa Flour 647 anti-CD45 (BioLegend, product 368537), W6/32 (1:500; ThermoFisher, product 14-9983-82), P-PI3K (1:1000; Cell Signaling Technologies, product 4228T), PI3K (1:1000; Cell Signaling Technologies, product 4292S), Histone H3 (Cell Signaling Technologies, product 4499T), B-Tubulin (Cell Signaling Technologies, product 2128S), anti-rabbit IgG HRP-linked (Cell Signaling Technologies, product 7074S), goat anti-mouse Alexa Flour 488 (Invitrogen, product A11001).

### Western blotting

1 x 10^6^ wild-type, PLCγ2^-/-^, or ATGL^-/-^ THP-1 cells were activated with 50 nM PMA for 48 hours in 6 well plates. Zymosan coated beads were added for 5, 10, 30, or 60 minutes at 37°C. Cells were rinsed 3 times with PBS and all residual liquid was removed before the addition of 2x Laemmli buffer to lyse the cells. The samples were scraped to collect samples and DNA was destroyed through mechanical perturbation and water bath sonication. Collected samples were loaded on a 12% poly-acrylamide gel. Proteins were transferred onto nitrocellulose membranes (Cytiva, product 10600013) with 100V current for 1 hour in the presence of transfer buffer containing Tris-Glycine base and 20% methanol. Membranes were blocked with TBS-T containing 5% BSA for 1 hour. Staining with antibodies was carried out in TBS-T containing 3% BSA for 1 hour. After antibody staining the membrane was treated with SuperSignal West Pico PLUS Chemiluminescent Substrate (ThermoFisher, product 34577) and imaged on an ImageQuant LAS400 (GE Healthcare) under the chemiluminescence setting.

### Quantification and statistical analysis

All statistical analysis was carried out using the R statistical software (version 4.3.2)^50^. To determine statistical significance multiple unpaired t-tests were performed using the pairwise_t_test function from the package rstatix^51^ and the Benjamini–Hochberg procedure for multiple comparison correction^52^.

### Binding assay

Binding assays were performed as previously described^28^ with the following modifications. 5 x 10^4^ wild-type, PLCγ2^-/-^, or ATGL^-/-^ THP-1 cells were activated with 50 nM PMA for 48 hours on coverslips in 24 well plates. After activation cells and beads were cooled at 10°C for 10 minutes before beads were added to cells at an MOI of 10. The samples were then spun at 277 x g for 3 minutes at 10°C to bring beads into contact with cells. Cells were transferred to a 37°C incubator with 5% CO_2_ for 20 minutes. After incubation cells were washed 3 times with ice-cold PBS before fixation with 4% formaldehyde for 30 minutes.

After fixation cells were stained with Concanavalin A (ConA; ThermoFisher, product C11252) and DAPI. For visualization and quantification (Figs 4A-B) images were acquired on a Nikon Spinning Disk confocal microscope (Nikon Yokogawa CSU-W1 Series Spinning Disk) with a Nikon Plan apochromat VC 60x Oil DIC N2 objective and 405 nm, 488 nm, and 561 nm lasers. Images were acquired in a 5 x 5 field and stitched together prior to analysis. Images were blinded and manually counted to identify beads in contact with, but not inside, cells.

### Receptor expression assay

To measure the surface expression and trafficking of surface receptors 5 x 10^4^ wild-type, PLCγ2^-/-^, or ATGL^-/-^ THP-1 cells were activated with 50 nM PMA for 48 hours on coverslips in 24 well plates. These cells were fixed for 30 minutes with 4% formaldehyde. Coverslips were treated with blocking buffer consisting of 5% normal goat serum (NGS; Rockland, product D204-00-0500), and either 0.1% Triton-X 100 or no permeabilizing agent for 1 hour at room temperature. Coverslips were then treated with primary antibodies diluted in blocking buffer with or without Triton-X 100 overnight at 4°C. Coverslips were washed 3 times with the proper blocking buffer and stained with the proper secondary antibody 1 hour at room temperature. Coverslips were again washed 3 times with the proper blocking buffer and stained with the conjugated anti-CD45 antibody for 1 hour at room temperature. These coverslips were washed 3 times in PBS and stained with DAPI for 5 minutes followed by fixation in 4% paraformaldehyde for 30 minutes. After fixation PBS was washed off coverslips with deionized water and the coverslips were mounted with ProLong Glass antifade mountant. For visualization and quantification (Figs 4C-D) images were acquired on a Nikon Spinning Disk confocal microscope (Nikon Yokogawa CSU-W1 Series Spinning Disk) with a Nikon Plan apochromat VC 60x Oil DIC N2 objective and 405 nm, 488 nm, 561 nm, and 640 nm lasers. Marker intensity was determined using the MeasureImageIntensity module in CellProfiler.

## Acknowledgements

We acknowledge expert technical assistance by staff in the Advanced Light Microscopy Core at the Jungers Center in the Department of Neurology at OHSU and a member of the University Shared Resources Program. This work is supported by R01AI141549 (F.G.T.), and 5T32GM142625 (A.G.).

